# Endothelial protein C receptor is an entry receptor for Langya virus

**DOI:** 10.64898/2026.11.04.763901

**Authors:** Young-Jun Park, Jaewon Song, Risako Gen, Cameron Stewart, Sanghyun Lee, David Veesler

**Author notes:** Equally contributions (Young-Jun Park, Jaewon Song, Risako Gen).

## Abstract

Langya virus (LayV) was recently discovered in infected individuals with febrile illness and classified as a *Parahenipavirus*, a group of poorly understood pathogens. LayV is distantly related to Nipah *Henipavirus* which causes recurring outbreaks with case fatality rates reaching up to 90%. No vaccines or therapeutics are approved for use in humans for any *Parahenipavirus* or *Henipavirus* despite their broad geographic distribution and pandemic potential. Here, we discovered that human endothelial protein C receptor (EPCR) is an entry receptor for LayV and that the tetrameric LayV attachment glycoprotein (G) binds EPCR avidly. We reveal the molecular basis of LayV G recognition of the EPCR receptor and provide a blueprint of the determinants of receptor ortholog usage. We found that human and shrew EPCR orthologs promote efficient cell entry of LayV and of the closely related Mojiang pseudoviruses indicating conserved receptor usage among this group of *Parahenipaviruses*. These results along with the detection of several *Parahenipaviruses* in shrews set these pathogens apart from *Henipaviruses*, which use Pteropid fruit bats as reservoir hosts. Our data support future surveillance efforts, by enabling prediction of EPCR utilization among newly discovered *Parahenipaviruses*, and the development and evaluation of countermeasures targeting this emerging group of human pathogens.

---

Paramyxoviruses are a diverse family that includes important human and animal pathogens. The discovery of Hendra virus (HeV) and of Nipah virus (NiV) in the 1990s led to the establishment of the *Henipavirus* viral genus of the Paramyxoviridae family^1,2^. NiV and HeV are characterized by a broad species tropism and can cause severe respiratory disease and encephalitis with high case fatality rates^3^. Recurrent zoonotic emergence from Pteropid fruit bats, with NiV outbreaks reported almost annually in Bangladesh and periodically in India, underscores the need for effective countermeasures. Subsequent sequencing and serological studies revealed the existence of additional *Henipaviruses* and their distribution over a much broader geographic distribution than initially appreciated with 2 billion people at risk of outbreaks^4–7^.

Langya virus (LayV) is distantly related to NiV and HeV and was described in 2022 after being found in febrile patients in China and in shrews^8^. Mojiang virus (MojV) is closely related to LayV and was identified in rats from an abandoned mine investigated after fatal pneumonia in mineworkers, although a causal association with illness was not established^9^. LayV and MojV are now classified in the *Parahenipavirus* genus which comprise members found on several continents, including the recent description of Camp Hill virus found in northern short-tailed shrews in Alabama(Parry et al. 2025; Vanmechelen et al. 2022; Imai et al. 2026; Caruso and Edwards 2023; Natasha et al. 2025; Chen et al. 2023).

Entry of *Henipaviruses* and *Parahenipaviruses* into cells rely on the concerted action of the attachment glycoprotein (G) and the fusion glycoprotein (F). G is a type II transmembrane attachment glycoprotein that assembles as homotetramers^14,15^, whereas F is a type I transmembrane glycoprotein that folds into metastable prefusion homotrimers^16^. Monoclonal antibodies with potent neutralizing activity have been identified against *Henipavirus* F and G and they conferred protection in small animal models and non-human primates^14,17–26^. The NiV/HeV G-directed m102.4 antibody was administered on a compassionate basis to several individuals with high-risk exposure to NiV or HeV. It has completed a phase 1 clinical trial and its efficacy in humans remains to be determined^27^. The humanized F-directed antibody hu1F5 provided greater protection than m102.4 in a nonhuman-primate study of NiV-Bangladesh infection and is advancing to clinical evaluation as MBP1F5^28^. Several NiV and HeV vaccine candidates based on G alone or F and G combined are under active development with some of them at the clinical evaluation stage^29–33^. However, NiV/HeV-directed monoclonal and polyclonal antibodies tested did not appreciably cross-react with the LayV or MojV F and G glycoproteins^15,34^. These antigenic differences raise concerns that vaccines developed against NiV and HeV may not provide cross-protection against *Parahenipaviruses*. Currently, no specific therapeutics or vaccines are approved for use in humans for any *Henipaviruses* or *Parahenipavirus*es.

*Henipaviruses* use the ephrin family of receptors to enter host cells and initiate infection. For instance, NiV and HeV can use the ephrin-B2 and ephrin-B3 receptors to infect hosts belonging to six distinct mammalian orders^35–38^. Upon ephrin-B2 or ephrin-B3 binding to G, F undergoes large-scale conformational changes to promote fusion of the viral and host membranes enabling delivery of the viral genome and initiation of infection^39,40^. Although several studies suggested that *Parahenipaviruses* do not use ephrins for engaging host cells, the identity of the host receptor and the tropism of these viruses have remained elusive^15,34,41^. The lack of known receptor has thus far limited the evaluation of countermeasures against these recently discovered pathogens for which no *in vitro* viral entry or replication assays or animal challenge models are available.

## EPCR is an entry receptor for LayV

To elucidate the mechanism of LayV entry into cells, we set out to discover cell surface proteins interacting with LayV G, which is expected to mediate viral attachment to target cells. We carried out a CRISPR activation (CRISPRa) screen using a *surface*ome library of 5,891 genes encoding for surface proteins with transmembrane domains, GPI-motifs, or that are associated with the plasma membrane. sgRNAs were transduced in human melanoma A375 cells stably expressing catalytically dead Cas9 to promote expression of these surfaceome genes individually. Following incubation with the LayV G ectodomain tetramer and staining with a fluorescently labelled antibody, we used flow cytometry to isolate positive cells for sgRNA sequencing. We found that the endothelial protein C receptor (EPCR encoded by the *PROCR* gene) was the most significantly enriched hit, followed by *GPC4*, *CR2* and *CD36* (**Fig 1A-B and Table S1**).

**Fig 1.**
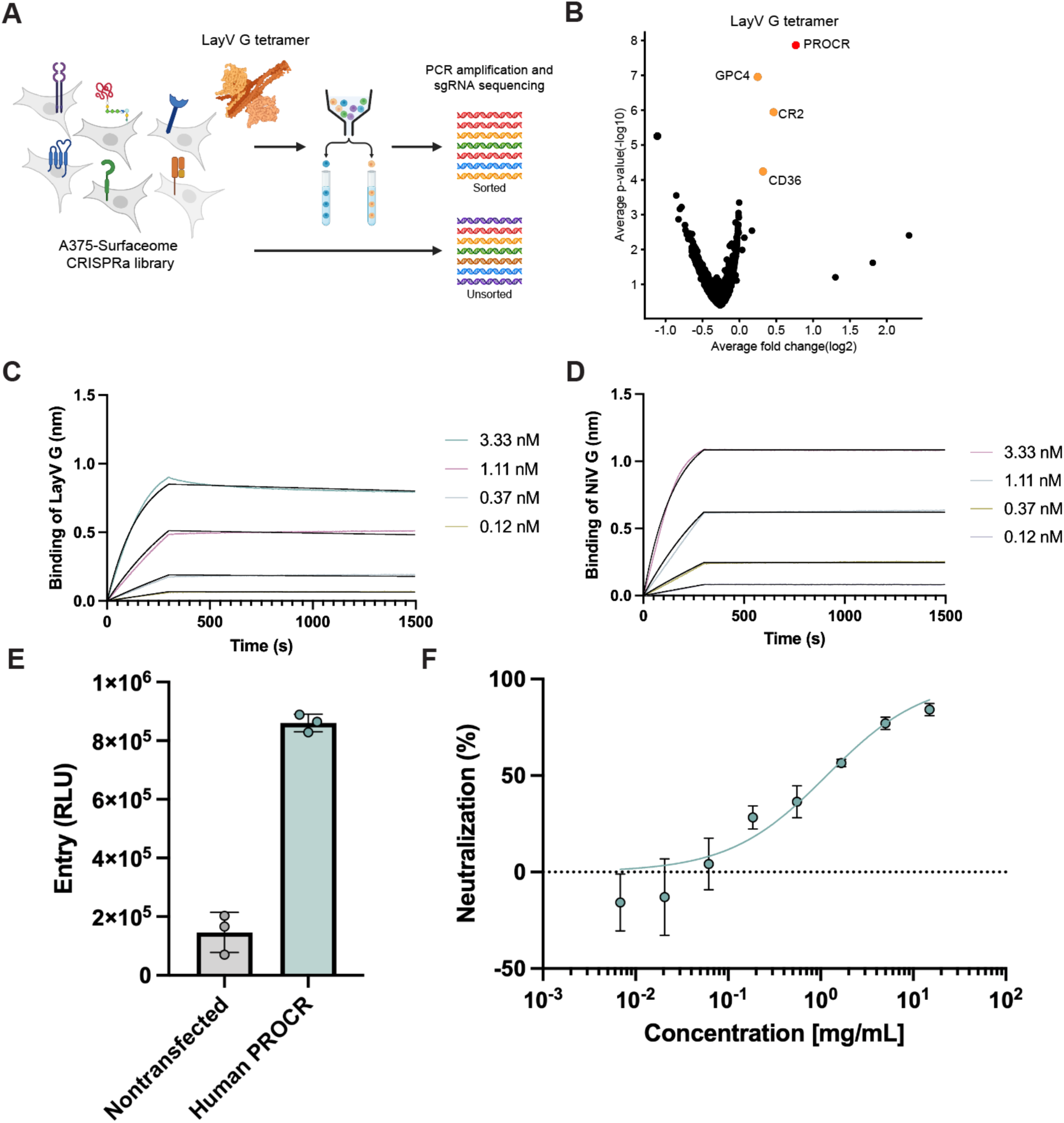
EPCR is an entry receptor for LayV. **A.** *Surface*ome CRISPR activation screening workflow. dCas9-expressing A375 cells were transduced with sgRNA library targeting 5,891 surface proteins, and cells binding to the LayV G ectodomain tetramer were sorted for sgRNA sequence analysis. **B**. Volcano plot showing sgRNAs enriched in cells binding to the LayV G ectodomain tetramer in the surfaceome CRISPRa screening shown in A. Fold change and p-value for each gene were calculated compared to the unsorted sample. *PROCR* is the gene encoding EPCR. **C-D**. Biolayer interferometry analysis of binding of the LayV G ectodomain tetramer to EPCR-Fc (C) or of the NiV G ectodomain tetramer to ephrin-B2-Fc (D). The ligands were immobilized at the surface of protein A biosensors. **E**. Entry of non-replicating, lentiviral particles pseudotyped with the LayV F/G glycoproteins in HEK293T cells transiently transfected or not with human EPCR. Each data point represents a technical replicate with error bars showing standard deviation. **F.** Soluble EPCR-mediated neutralization of lentiviral particles pseudotyped with the LayV F and G glycoproteins. Each data point represents the average of two technical replicates with error bars showing standard deviation.

To validate the results from the *surfaceome* screen, we first used CRISPRa to induce endogenous EPCR expression in the A375 and HeLa human cell lines with two sgRNAs individually targeting the *PROCR* promoter region. Both sgRNAs (sgPROCR #1 and #2) effectively induced cell surface expression of EPCR and substantially increased binding of LayV G compared to non-induced controls (**Fig S1A-B**). Subsequent ectopic expression in HEK293T cells of each of the four hits identified in the *surfaceome* CRIPRa screen showed that EPCR led to a marked enhancement of LayV G binding relative to other candidates evaluated or to untransfected controls (**Fig S1C-E**). EPCR is a type I transmembrane protein with an MHC class I-like ectodomain involved in control of blood coagulation and expressed at the surface of endothelial cells as well as several other cell types(Laszik et al. 1997; Wang et al. 2007; Galligan et al. 2001; Bretschneider et al. 2007; Oganesyan et al. 2002).

We then aimed to determine whether LayV G engages EPCR directly, rather than through additional cell surface components. Using biolayer interferometry, we found that LayV G bound to EPCR but not ephrin-B2 whereas NiV G bound ephrin-B2 but not EPCR, confirming engagement of distinct cell surface molecules by these two distantly related viruses. The LayV G and NiV G ectodomain tetramers respectively bound to EPCR and ephrin-B2 with picomolar avidities due to the presence of four receptor-binding sites on each G tetramer (**Fig 1C-D**). To determine the functional relevance of LayV G binding to EPCR, we evaluated the influence of transient transfection of HEK293T cells with membrane-anchored EPCR on entry of a non-replicating, lentivirus pseudotyped with the LayV F and G glycoproteins. Transfection of human EPCR enhanced the susceptibility of cells to LayV F/G-mediated entry and soluble EPCR could inhibit entry in a concentration-dependent manner (**Fig 1F-G and Fig S1F-G**). Collectively, these results identify EPCR as a *bona fide* entry receptor promoting LayV F/G-mediated viral entry.

## Molecular basis of LayV G recognition of the EPCR receptor

To decipher the molecular basis of LayV engagement of its host receptor, we characterized the LayV G ectodomain tetramer bound to the EPCR ectodomain using cryoEM. Data processing revealed the presence of homogeneous LayV G homotetramers with the head domains stoichiometrically bound to EPCR. In the structure, the four LayV G head domains are positioned against the central 4-helix bundle stalk, yielding a C4 symmetrical structure at 2.9Å resolution (**Fig 2A-B, Figure S2 and Table S2**). LayV G uses the lateral side of its β-propeller head domain to bind EPCR, thereby burying an average ∼670Å^2^ of the surface of each binding partner. LayV G inserts residue F336 into the groove formed by the MHC I-like EPCR fold contacting receptor residues and the aliphatic tail of the phosphatidyl-ethanolamine occupying the cleft in all EPCR structures (**Fig 2C**). Furthermore, a salt bridge between R98_EPCR_ and D315_LayVG_ and a hydrogen bond between Q92_EPCR_ and the Y338_LayVG_ backbone carbonyl further reinforce these interactions along the ridge of the EPCR cleft (**Fig 2C-D**). The importance of these interactions is underscored by the fact that the LayV G R_332_ITGFAYL_339_ protrusion accounts for approximately 60% of the LayV G surface buried at the interface with EPCR.

**Figure 2.**
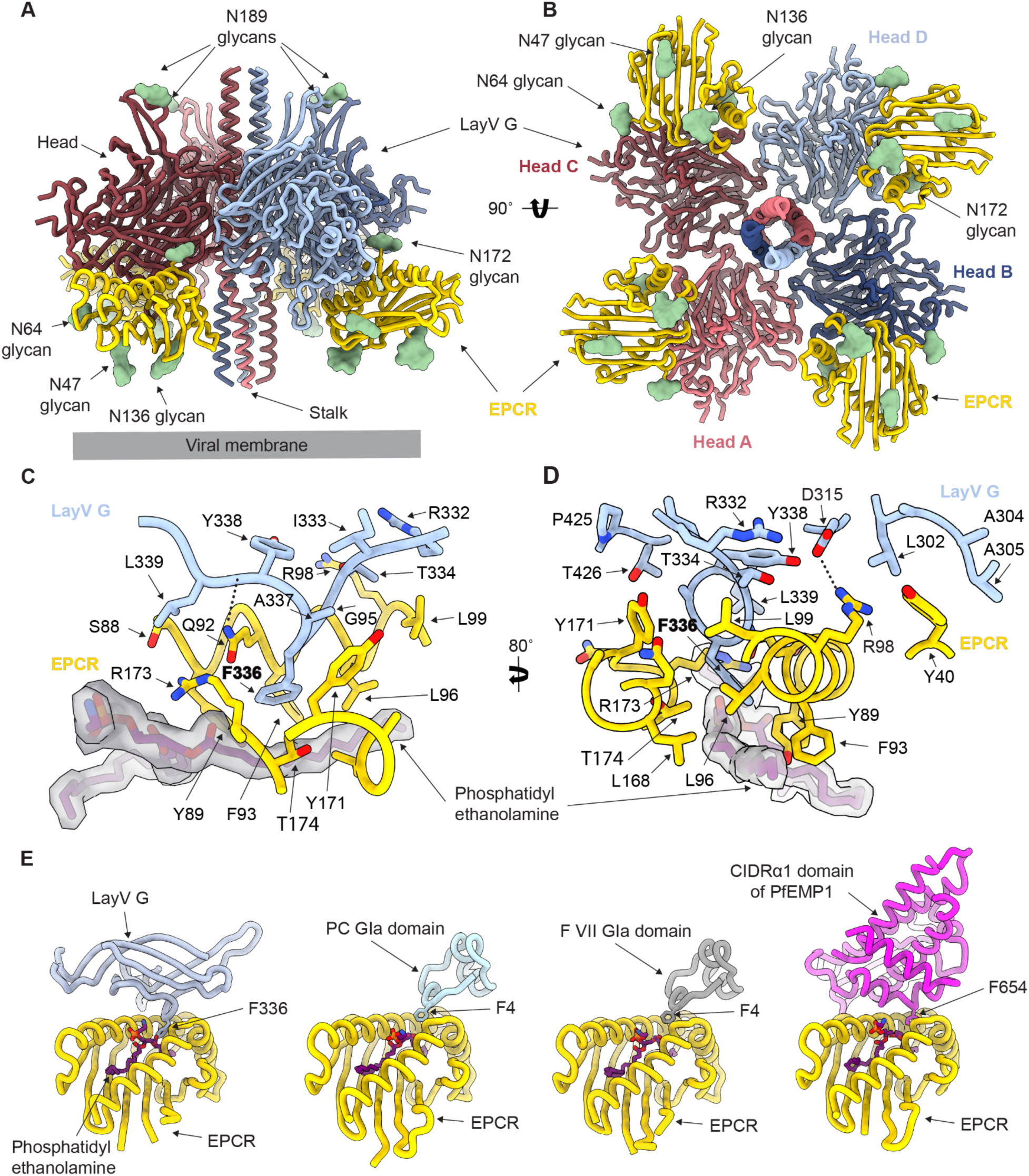
Molecular basis of LayV G engagement of the human cell surface receptor EPCRgg. **A-B**. CryoEM structure of the human EPCR receptor bound LayV G (oP4H-LayV G_SM_) tetramer rendered as ribbons shown in two orthogonal orientations. N-linked glycosylation sites are rendered as green surfaces. **C-D**. Zoomed-in view of the interactions between LayV G head domain and human EPCR receptor with select side chains shown as sticks. The EPCR phosphatidyethanolamine is rendered as sticks with the cryoEM map as semi-transparent surface. **E**. Comparison of the EPCR binding modes of the receptor binding domain of LayV G, the Protein C gamma-carboxyglutamic acid (Gla) domain (PDB: 1LQV), the Factor VII Gla domain (PDB: 9TJX), and the CIDRα1 domain of *Plasmodium falciparum* erythrocyte membrane protein 1 (PfEMP1) (PDB: 4V3E). EPCR residues are numbered according to the UniProt Q9UNN8 precursor, including the 17-residue signal peptide, which corresponds to a 17 residue offset with the numbering used in previously deposited EPCR structures.

Comparison of the EPCR-bound LayV G structure with the EPCR-bound protein C Gla domain^42^, coagulation factor VII Gla domain^43^ and with the Plasmodium falciparum erythrocyte membrane protein 1 (PfEMP1) structures^44,45^ reveals a conserved recognition strategy (**Fig 2E**). In all four cases, a Phe residue from the binding partner protrudes in the EPCR cleft to recognize overlapping sites, indicating convergence of molecular solutions adopted by multiple, unrelated EPCR-binding partners.

The overall LayV G structure adopts a conformation similar to apo LayV G (PDB 8TVI) to which it can be superimposed with an RMSD of 0.44 Å over 2,036 aligned Cα positions (**Fig S3A**). In this conformational state, the receptor-binding motif on each LayV G head is oriented towards the viral membrane instead of the host membrane (**Fig 2A-B and Fig S3B)**. This observation is reminiscent of the NiV G architecture in which 3 out of 4 ephrinB2 receptor-binding motifs within a tetramer are oriented towards the viral membrane, indicative of conformational changes associated with the entry process. However, we note that ephrinB2 and EPCR recognize distinct surfaces on the NiV and LayV G heads, respectively (**Fig S3C-D**). These results provide a blueprint of the interactions enabling EPCR to promote LayV F/G-mediated entry into cells.

## LayV G and MojV use a broad spectrum of EPCR orthologs

To understand which other related viruses may use EPCR as an entry receptor, we examined the receptor-binding motif across *Parahenipavirus* G glycoproteins. EPCR-interacting G residues are invariant across the 11 sequenced (8 unique) LayV genomes, irrespective of their human or *Crocidura* origin(Natasha et al. 2025; Zhang et al. 2022). Furthermore, an aromatic residue is found at the position equivalent to LayV G F336 in the three *Parahenipaviruses* closest to LayV: MojV G (Y336), Wenzhou shrew henipavirus 1 G (F362) and *Crocidura olivieri* henipavirus 1 G (Y338) which respectively share 86%, 69% and 68% sequence identity with LayV G(Chen et al. 2023) (**Fig 3A**). This observation along with the conservation of other surrounding residues (D315, R332, G335, A337, Y338 and L339), suggests shared receptor usage among these four viruses (**Fig 3A**). However, substitutions present in this region for other *Parahenipaviruses* harboring more distant G glycoproteins, including mutations at the position equivalent to the LayV G F336 residue by Arg, Lys, Ser, Glu or Asn are likely incompatible with EPCR utilization (**Fig 3A**). To test this hypothesis, we evaluated entry of a non-replicating lentivirus pseudotyped with the MojV F and G glycoproteins into HEK293T cells transiently transfected with human EPCR. We found that human EPCR rendered cells susceptible to MojV F/G-mediated entry, establishing it as an entry receptor for this *Parahenipavirus* as well (**Fig 3B**). Collectively, these findings suggest that LayV, MojV, Wenzhou shrew henipavirus 1 and C*rocidura olivieri* henipavirus 1 form a subgroup of *Parahenipaviruses* that have evolved to use EPCR as an entry receptor.

**Figure 3.**
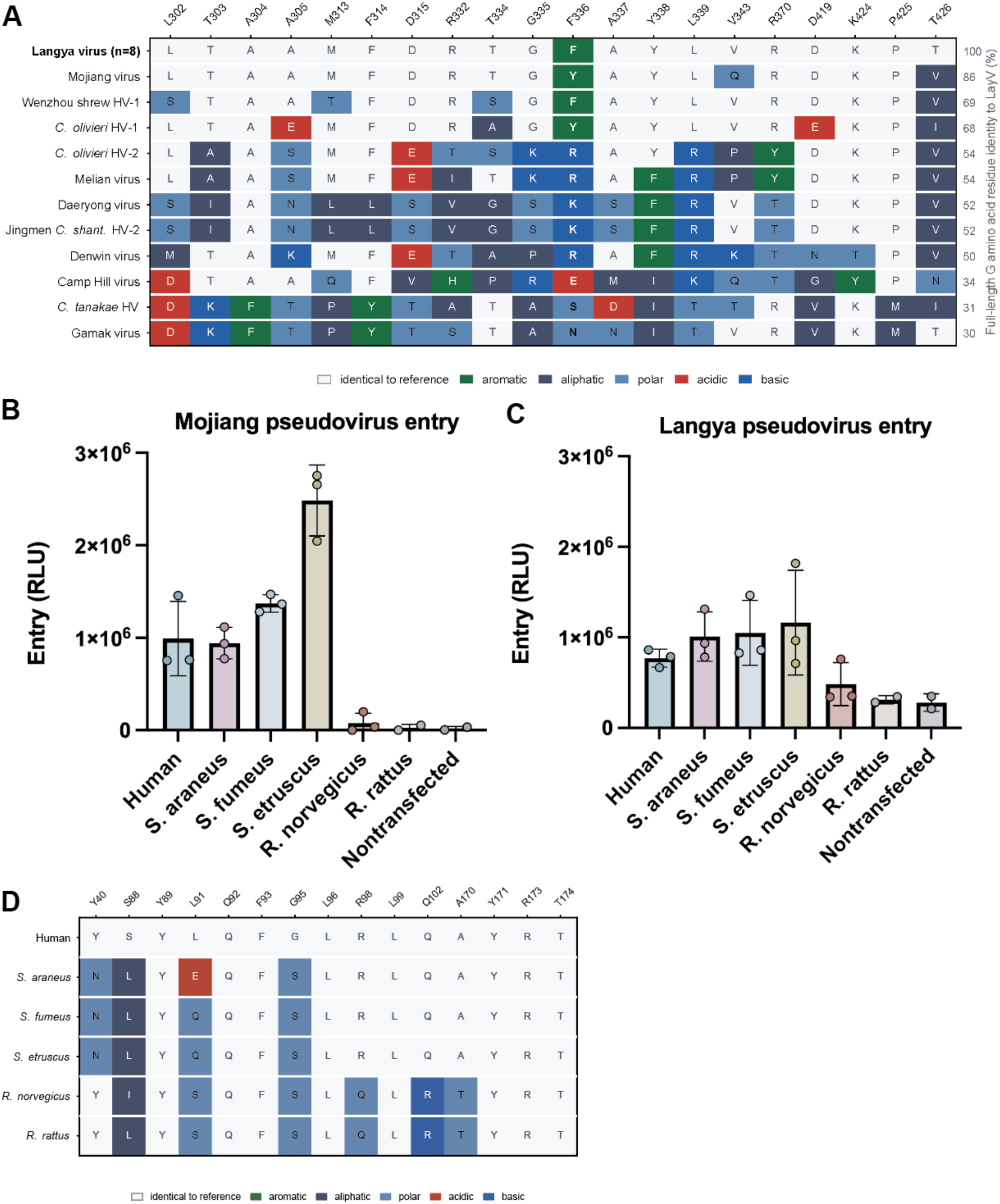
LayV G and other *Parahenipaviruses* use a broad spectrum of EPCR orthologs. **A**. Sequence conservation of *Parahenipavirus* G residues binding to EPCR. Accession numbers: Langya virus, UUV47241.1, UUV47206.1, UUV47213.1, XLS83347.1, XLS83377.1, XLS83387.1, XLS83357.1, XLS83367.1 (n = 8 distinct G sequences out of 11 genomes); Mojiang virus, YP_009094095.1; Wenzhou shrew henipavirus 1, WPV62327.1; Crocidura olivieri henipavirus 1, XOT37559.1; Crocidura olivieri henipavirus 2, XOT37566.1; Melian virus, YP_013033284.1; Daeryong virus, YP_013025921.1; Jingmen Crocidura shantungensis henipavirus 2, UOX72991.1; Denwin virus, YP_013033267.1; Camp Hill virus, XJU75811.1; Crocidura tanakae henipavirus, WIU81493.1; Gamak virus, QYO90518.1. **B-C**. Assessment of MojV F/G and LayVF/G-mediated entry of non-replicating, lentiviral pseudotyped virus entry into HEK293T cells transiently transfected with the *Sorex araneus* (XP_004612598.1), *Sorex fumeus* (XP_055977907.1), *Suncus etruscus* (XP_049635309.1), *Rattus norvegicus* (Q4V8I1), *Rattus rattus* (XP_032760446.1), or human (Q9UNN8) EPCR orthologs. Each dot represents a biological replicate using different batches of pseudovirus and transfected cells with error bars showing standard deviation. **D.** Sequence conservation of EPCR residues interacting with G.

Given the detection of LayV RNA in shrews and of MojV RNA in rodents, we set out to investigate the receptor ortholog usage of these viruses to further our understanding of receptor compatibility and spillover potential. We therefore evaluated entry of LayV and MojV pseudoviruses into HEK293T cells transiently transfected with *Sorex araneus*, *Sorex fumeus*, *Suncus etruscus*, *Rattus norvegicus, Rattus rattus,* or human EPCR orthologs. We found that human, *S. araneus*, *S. fumeus*, and *S. etruscus* EPCR orthologs promoted high entry levels of LayV and MojV pseudoviruses, whereas *R. norvegicus*, and *R. rattus* EPCR orthologs enabled low entry of LayV and no detectable entry of MojV (**Fig 3B-C, Figure S4**). These results reflect the high sequence conservation of the key amino acid residues lining and directly surrounding the LayV G F336-interacting pocket among the human and shrew EPCR orthologs evaluated here (**Fig 3D**). There are 3 mutations at the LayV G-binding interface setting the rodent EPCR orthologs apart from the other EPCR tested in our panel: R98Q, Q102R and A170T. The R98Q substitution would abrogate the salt bridge formed with LayV G D315 in our structure and may explain the weak utilization of rodent EPCRs (**Fig 2D**). Collectively, our data indicate that multiple *Parahenipaviruses* related to LayV utilize several EPCR orthologues as host cell entry receptors.

## Discussion

EPCR is a transmembrane glycoprotein with an MHC class I-like ectodomain^46^ mainly expressed on vascular endothelial cells along with selected non-endothelial populations, including alveolar epithelial cells, monocytes and vascular smooth muscle cells^47–50^ as well as cultured rat cortical and hippocampal neurons^51^. A major physiological role of EPCR is to negatively regulate the blood coagulation cascade by binding protein C and enhancing its activation by the thrombin-thrombomodulin complex, thereby promoting anti-coagulant activity^52^. EPCR also supports activated protein C-dependent cytoprotective signaling through protease activated receptor 1^53^ and it also contributes to the pathogenesis of severe *Plasmodium falciparum* malaria by binding to PfEMP1 displayed on the surface of infected erythrocytes^44,54^. Given that LayV G engages the same EPCR cleft as the one used by protein C, factor VIIa and PfEMP1, LayV G and any of these ligands cannot occupy EPCR simultaneously. LayV was initially identified and isolated from a patient’s throat-swab, and laboratory abnormalities reported in infected patients include leukopenia, thrombocytopenia, and impaired hepatic and renal function ^8^. LayV G recognition of EPCR would therefore be expected to block protein C binding to the same receptor and reduce the EPCR-dependent enhancement of protein C activation by the thrombin-thrombomodulin complex. Whereas the haematological abnormalities reported in infected individuals have not been assigned to a defined infected cell type, this relationship offers a testable hypothesis for them and a mechanism by which *Parahenipavirus* attachment could perturb endothelial anticoagulant function. The distantly related NiV and HeV use ephrin-B2 and ephrin-B3 as entry receptors(Negrete et al. 2005; Bonaparte et al. 2005) and can cause respiratory disease and encephalitis(Eaton et al. 2006; Goh et al. 2000). Future work will be needed to determine if LayV shares the cell and tissue tropism as well as disease pathology established for these highly lethal *Henipaviruses* despite using distinct receptors.

The EPCR-bound LayV G structure presented here adopts a conformation indistinguishable from the apo LayV G, with all four head domains positioned against the central 4-helix stalk, irrespective of the presence or absence of its transmembrane domain^15,55^. This architecture is distinct from that observed in the NiV G tetrameric structure which adopts an asymmetric conformation with two β-propeller heads positioned against the central 4-helix stalk and the other two heads positioned distally from the viral membrane and with opposite orientation to each other^14^. As a result, each NiV G tetramer orients one out of four receptor-binding motifs towards the host membrane whereas LayV G positions all four of them towards the viral membrane. Prior work on NiV and HeV suggested that receptor engagement induces G conformational changes leading to exposure of the neck domain, which is located in between the stalk and heads, to promote interactions with F and initiation of membrane fusion^39^. The absence of detectable EPCR-induced LayV G conformational changes raises the question as to whether receptor-binding alone is sufficient to induce this cascade of structural rearrangements or if additional biochemical cues are needed to initiate the entry process.

The detection of LayV in tens of human individuals indicates that *Parahenipaviruses* represent a zoonotic risk and that surveillance studies should monitor the known reservoir hosts as well as other animal species for outbreak preparedness. Indeed, we reveal here that LayV G binds an EPCR crevice which was also independently selected by the Plasmodium falciparum PfEMP1, with limited sequence variability due to its role in interacting with protein C and factor VIIa. Consequently, LayV, MojV and possibly closely related *Parahenipaviruses* can utilize EPCR orthologs from other mammalian species which are known hosts, including several shrew species. Broad receptor species tropism would expand the range of possible hosts for these pathogens and their risk of spreading, evolving and spilling over to humans. Moreover, our data provide a blueprint for predicting EPCR receptor utilization as *Parahenipaviruses* are discovered solely based on the G glycoprotein sequence. We note that productive infections not only result from cell entry but also require viruses to successfully thwart host innate immune responses and subvert cellular machineries for production of progeny virions. Future studies should integrate all these aspects to assess potential spillover and transmission risks.

The lack of cross-reactivity with LayV or MojV F and G glycoproteins of NiV or HeV-elicited monoclonal and serum polyclonal antibodies tested underscores the importance to develop countermeasures specifically for these *Parahenipaviruses* to enable rapid response in case of future emergence^15,34^. The discovery of EPCR as the LayV and MojV receptor paves the way to evaluate vaccines, monoclonal antibodies and other inhibitors as it enabled us to develop a non-replicating pseudovirus-based surrogate entry assays and will support *in vitro* viral propagation. Furthermore, these results will allow the future development of animal challenge models which will support *in vivo* assessment of the protective efficacy conferred by newly developed *Parahenipaviruses*-directed countermeasures and help understand pathogenesis of these poorly characterized pathogens.

## Acknowledgements

This work was supported in part by the NIH/NIAID under grants 1U19AI181881 and 75N93022C00036 to D.V., R01AI183155 and P20GM156688 to S.L., an Investigators in the Pathogenesis of Infectious Disease Awards from the Burroughs Wellcome Fund to D.V., the University of Washington Arnold and Mabel Beckman Cryo-EM Center, the Smith Family Awards Program for Excellence in Biomedical Research to S.L., and the Charles H. Hood Foundation Child Health Research Award to S.L. D.V. is an investigator of the Howard Hughes Medical Institute, and the Hans Neurath Endowed Chair in Biochemistry at the University of Washington.

## Contributions

JS RG, SL and DV designed the project. JS performed the CRIPRa screen and flow cytometry analysis. RG carried out pseudovirus entry assays. CS produced recombinant glycoproteins and carried out SPR analysis. YJP carried out the cryoEM work. All authors analyzed the data. DV wrote the manuscript with input from all authors. SL and DV supervised the project and acquired funding.

## Methods

### Cell lines and culture conditions

CHO-S (R80007) cells were used for EPCR ectodomain protein production by Genscript. For other recombinant protein production, Expi293F (A14527) used for soluble protein production was obtained from Thermo Fisher Scientific, cultured in Expi293 Expression Medium in suspension at 130 RPM, 37 °C and 8% CO_2_.

HEK-293T (ATCC) was grown in Dulbecco’s Modified Eagle Medium (DMEM) supplemented with 10% fetal bovine serum (FBS) and 1% penicillin–streptomycin–l-glutamine (PS) at 37 °C and 5% CO_2_ for pseudovirus production and entry assays.

For CRISPR and flow cytometry, HEK293T (CRL-3216), HeLa (CCL-2), and A375 (CRL-1619) cells were purchased from ATCC. Cells were cultured using DMEM (Gibco, 11995-081) supplemented with 10% FBS (Biowest, S1620) and 25 mM HEPES (Gibco, 15630-080) at 37°C, 5% CO_2_. 0.05% trypsin-EDTA with phenol red (Gibco, 25300-120) was used to detach cells. For HeLa and A375 cells, 5 µg/mL blasticidin (Gibco, A1113903) and 0.7∼1 µg/mL puromycin (Gibco, A1113803) were added as appropriate.

### Plasmids and vectors

For expression of the stabilized LayV G ectodomain tetramer (designated LayV G_SM_), the DNA sequence for the LayV G ectodomain (Genbank ID:UUV47206.1, residues 72-624) containing F89L, G98T, G117A, F126V, and Y130L stabilization mutations and N-terminal secretion signal peptide (MTRLTVLALLAGLLASSRA) derived from human azurocidin followed by His-tag (SGPHHHHHHHH) with flexible linker (GSSP) was human codon-optimized, synthesized, and inserted into the pCMV vector^15^. For CryoEM structure determination, a designed tetramerization domain (oP4H) sequence (QAEELKAIKEELKAIKEELKAIKEELKAI) was fused at the N-terminal of LayV G_SM_ as described in previous studies (designated oP4H-LayV G_SM_)^15^. The Nipah G (Genbank: NP_112027.1) ectodomain residues 70-602 were cloned into a pCMV construct with N-terminal 6XHis tag and a Mu-phosphatase signal peptide, with stabilizing stalk mutations. For recombinant EPCR protein, human EPCR ectodomain (Uniprot ID:Q9UNN8, residues 1-210) sequences with C-terminal Avi-tag (GLNDIFEAQKIEWHE) linked by linker sequences (GSS) followed by a hexahistidine-tag was human codon-optimized, synthesized, and inserted into pcDNA3.4 vector by Genscript. Human ephrinB2 (Uniprot ID: P52799) ectodomain residues 1-229 were cloned into a pCMV construct fused to human Fc.

For surface expression of EPCR (*PROCR*), GPC4, CR2, and CD36 proteins, the coding region of the target protein was synthesized and cloned into pcDNA3.1(+) plasmid with C-terminal Flag tag (Genscript).

For pseudovirus production, human EPCR membrane-anchored (Genbank ID:Q9UNN8, residues 1-238) sequences with C-terminal GSSG linker sequences followed by HA Tag (YPYDVPDYA) was was human codon-optimized, synthesized, and inserted into pcDNA3.4 vector by Genscript. Full-length Langya F (Genbank ID: UUV47240.1) with a C-terminal Flag tag (DYKDDDDK), and Langya G (Genbank ID: UUV47241.1) with an N-terminal Flag tag, were both cloned into pCMV vectors. Full length Mojiang F (Genbank ID: YP_009094094) with a C-terminal S tag (KETAAAKFERQHMDS) and Mojiang G (Genbank ID: YP_009094095) were cloned into pCDNA3.1(+) vectors.

### Lentivirus constructs and stable cell lines generation

For CRISPR activation, pLenti-dCas9-VP64-Blast (Addgene, 61425) was used to express catalytically dead Cas9, and pXPR_502 (Addgene, 96923) was used to express individual sgRNAs targeting PROCR (sgPROCR #1: AGAAGTGAGGACCGTCTGGG; sgPROCR #2: GTGTCTCATTTCCTACTAGG). HEK293T cells were transfected with a lentiviral plasmid, pMD2.G (Addgene, 12259) and psPAX2 (Addgene, 12260), using TransIT-LT1 (Mirus Bio, MIR 2300). After replacing the culture media at 6 hours post transfection, the supernatant was harvested and filtered using a 0.45 µm filter (Millipore) at 36 hours post transfection. For transduction, A375 or HeLa cells were transduced with 1 mL of lentiviral stock by centrifuging at 1200 x g for 90 minutes at 30°C with 1 µg/mL polybrene. At 18 hours post transduction, the culture media was replaced and cells were selected with appropriate drugs for 7 days.

### Surfaceome-focused CRISPR activation screen

Surfaceome-focused CRISPR activation screen was performed as previously described^56^ with minor modifications. Briefly, A375-dCas cells expressing CRISPR activation sgRNAs targeting 5,891 genes were incubated with 50 µg/mL LayV G or NiV G-His tetramer protein in FACS buffer (1x phosphate buffered saline (PBS) supplemented with 2% FBS and 1 mM EDTA) for 30 min at 4°C, then fixed with 4% formaldehyde for 10 min at room temperature. After washing twice with FACS buffer, cells were sequentially stained with anti-His (Abcam, ab18184) and PE anti-mouse IgG antibody (Biolegend, 405307). Cells were sorted using Cytek Aurora CS (Cytek Biosciences) and the top ∼3% (fluorescence intensity) of the PE-positive cells were isolated.

Genomic DNA was extracted from the sorted and unsorted cells using QIAamp DNA Mini or Maxi kit (Qiagen, 51104, 51192). After PCR amplification using Titanium Taq DNA polymerase (Takara, 639209), P5 stagger primer mix (0.5 µM) and uniquely barcoded P7 primer (0.5 µM), amplicon DNA samples were purified with AMPure XP beads (Beckman Coulter, MSPP-A63880). Next-generation sequencing was conducted using NextSeq 2000 sequencer (Illumina), and reads were demultiplexed and mapped to each sgRNA using Library Aligner (https://gitlab.com/buchserlab/library-aligner). sgRNA enrichment in each sorted population was quantified as the log2 fold change versus the unsorted population, with the p-values derived from the hypergeometric distribution.

The analyzed screening results are provided as Table S2 and the raw FASTQ files were deposited in the NCBI Sequence Read Archive Bioproject PRJNA1556238.

### Flow cytometry

Langya G protein binding to cells expressing EPCR was performed using CRISPRa cell lines in A375 and HeLa, or cDNA-transfected HEK293T cells. For CRISPRa cell lines, A375- or HeLa-dCas cells were transduced with individual activating sgRNAs targeting PROCR. For cDNA-transfection, HEK293T cells were seeded into 6-well plate and incubated for 24 hours, then subsequently transfected with pcDNA3.1 plasmids bearing the receptor gene with the C-terminal Flag tag using TransIT-LT1 (Mirus Bio, MIR 2300). One day after transfection, receptor expressing cells were subjected to flow cytometry binding assay. Cells were incubated with 100 µg/mL of LayV G-His tetramer protein in HBSS containing 2% FBS for 30 minutes at 4°C, then fixed with 4% paraformaldehyde for 10 minutes at room temperature. Cells were washed twice with FACS buffer (1x PBS containing 2% FBS and 1 mM EDTA) and stained with anti-His antibody (Abcam, ab18184) and PE anti-mouse IgG (Biolegend, 405307) in FACS buffer for 30 minutes at 4°C. To measure surface EPCR expression, cells were first stained with a rat monoclonal anti-EPCR antibody (clone RCR-401; Biolegend, 351902) in FACS buffer, then stained with AF488-conjugated goat polyclonal anti-rat IgG (Thermo Fisher Scientific, A-11006) after washing. Cells were analyzed using Cytek Aurora spectral analyzer (Cytek Biosciences) and data was analyzed with FlowJo software.

### Protein expression and purification

Soluble LayV G ectodomain (residues 72-624) used for Cryo-EM or BLI assays was produced in Expi293F cells. Following the manufacturers protocols Expi293F cells were grown to a density of 3 x 10^6^ cells/mL and transfected using the ExpiFectamine293 Transfection Kit (ThermoFisher Scientific). The expression continued for 5 days post transfection, with the cells staying at 37°C, 8% CO_2_ and constant rotation at 130 RPM during the course of expression. On the final day the cell supernatant was clarified via centrifugation at 6000xg then purified using HisPur Ni-NTA Resin (Thermo) packed in gravity flow columns (Marvelgent Biosciences). The resin was washed with 20 column volumes(CVs) of 50 mM Tris, 150 mM NaCl, 10 mM imidazole pH 7.5 before elution with 10 CVs of 50 mM Tris, 150 mM NaCl, 500 mM imidazole pH 7.5. Shortly after affinity purification the LayV G ectodomain was further purified via size exclusion chromatography in a Superose 6 Increase 10/300 GL (Cytiva) into a final storage buffer of 50 mM Tris, 150 mM NaCl pH 7.5. Fractions containing LayV G protein were pooled then promptly flash frozen to preserve for future use.

EPCR ectodomain protein was produced using CHO-S cells and was purified using Ni Magnetic beads (Genscript, L007764) in PBS buffer pH 7.2 by Genscript. Fractions containing EPCR protein were pooled then flash frozen to preserve for future use.

Human ephrinB2-Fc dimer was produced in Expi293F cells grown in suspension using Expi293 Expression Medium (Thermo Fisher) at 37°C in a humidified 8% CO2 incubator rotating at 130 rpm. The cultures were transfected using ExpiFectamine™ 293 Transfection Kit for Expi293 cells (Thermo Fisher) with cells grown to a density of 3 million cells per mL and cultivated for 5 days. The supernatants were harvested, and proteins were purified from clarified supernatants using a protein A affinity column. Proteins were further purified via size exclusion chromatography on a Superdex200 Increase 10/300 GL (Cytiva), and stored in 50 mM Tris, 150 mM NaCl at pH 8.0.

The NiV G ectodomain was produced in Expi293F cells grown in suspension using Expi293 Expression Medium (Thermo Fisher) at 37°C in a humidified 8% CO2 incubator rotating at 130 rpm. The cultures were transfected using ExpiFectamine 293 Transfection Kit for Expi293 cells (Thermo Fisher) with cells grown to a density of 3 million cells per mL and cultivated for 5 days. The supernatants were harvested, and proteins were purified from clarified supernatants using a 1mL Cobalt affinity column (Takara). Proteins were further purified via size exclusion chromatography on a Superose 6 Increase 10/300 GL (Cytiva), and stored in 50 mM Tris, 150 mM NaCl at pH 8.0.

### Pseudotyped virus production

A single-cycle (replication-incompetent) HIV-1 envelope-deficient backbone vector containing a luciferase reporter was pseudotyped with Langya or Mojiang F and Langya or Mojiang G glycoproteins. Briefly, 15-20 million HEK293T (ATCC, CRL-11268) were seeded in T175 flasks in DMEM (ThermoFisher, Gibco 11-965-118) supplemented with 10% FBS (Cytiva, Hyclone SH3039603HI), and incubated overnight at 37°C with 5% CO_2_. The following day, a mixture of 1.5 mL DMEM, 15 ug pHage-CMV-Luc2-IRES-ZsGreen-W, 3.3 ug pRC-CMV-ReV1b, 3.3 ug HDM-tat1b, 3.3 ug HDM-Hgpm2, 5.1 ug Langya/Mojiang F and 5.1 ug Langya/Mojiang G was prepared before adding 87.8 uL of Lipofectamine 2000 (ThermoFisher, Invitrogen 11668500), and incubated at room temperature for 5-15 minutes. The transfection mixture was subsequently added to the T175 flasks containing 20mL of new DMEM supplemented with 10% FBS. Flasks were then returned to incubate at 37°C with 5% CO_2_ overnight. After ∼24hrs, 200uL sodium butyrate (from 500mM stock prepared in filtered MilliQ water) was added to the flasks to achieve a final concentration of 5mM in 20mL of media. Flasks were returned to incubate at 37°C with 5% CO_2_ overnight. After ∼24hrs, supernatant was collected from each flask, spun down at 2500xg for 10 min, filtered through a 0.45 um membrane, and concentrated 20x using a 100 kDa cut-off membrane (Millipore Sigma, Amicon UFC903024). Pseudoviruses were then aliquoted and frozen at −80°C until use.

### Pseudovirus entry and neutralization assay

For LayV/MojV pseudovirus entry assays, HEK293T cells were seeded into T75 flasks containing DMEM (ThermoFisher, Gibco 11-965-118) supplemented with 10% FBS (Cytiva, Hyclone SH3039603HI) and incubated overnight at 37°C with 5% CO_2_. The following day, when cells reached 90% confluency, a mixture of 10.5 ug EPCR full-length plasmid, 740 uL DMEM, and 26.4 uL of Lipofectamine 2000 (ThermoFisher, Invitrogen 11668500) were added to the cells after 5-15 minutes of incubation. After 4-5hrs, cells were detached, counted and reseeded at ∼30,000 cells per well in poly-lysine-coated 96 well plates. Cell plates were returned to incubate overnight at 37°C with 5% CO_2_. The next day, a half-area 96-well plate was prepared with 1:4 starting dilution of pseudovirus and diluted serially at a 1:2 dilution. The media was removed from transfected HEK293T cells and pseudovirus was transferred to the cells along with 5 uL of 50 ug/mL polybrene (Sigma-Aldrich), and incubated for 3 h at 37°C with 5% CO_2_ before adding 40 uL of 10% FBS and 1% PenStrep (100 I.U./mL penicillin and 100 mg streptomycin) containing DMEM. Following 48 h incubation, 40 uL of One-GloEX (Promega E8150) was added to the cells, placed on a shaker, and incubated in the dark for 3 min. Plates were then read on an Agilent BioTek Neo2 plate reader at maximum gain of 255, and relative luciferase units were plotted in Prism (GraphPad). At least two biological replicates with two distinct batches of pseudovirus were conducted.

For LayV/MojV neutralization assays, cells were transfected and seeded in the same way as stated above. The next day, a half-area 96-well plate was prepared with soluble EPCR at a starting final concentration of 15-22.5 mg/mL, serially diluted 1:2 in DMEM in 22 uL final volume. 22 uL of pseudovirus was then added to each well and incubated at 37°C for 30-45 min. The media was removed from transfected HEK293T cells and 40 uL of the EPCR/pseudovirus mixture was transferred to the cells and incubated for 3 h at 37°C with 5% CO_2_ before supplementing with an additional 40 uL of 10% FBS and 1% PenStrep containing DMEM. Following 48 h incubation, 40 uL of One-GloEX (Promega E8150) was added to the cells, placed on a shaker and incubated in the dark for 3 min. Plates were then read on an Agilent BioTek Neo2 plate reader. Relative luciferase units were plotted and normalized in Prism (GraphPad) using a zero value of cells alone and a 100% value of 1:2 virus alone. Nonlinear regression of log(inhibitor) vs. normalized response was used to determine EC50 values from curve fits. At least two biological replicates with two distinct batches of pseudovirus were conducted.

### Western blot analysis

For EPCR expression western blot analysis, transfected cells in a 96-well plate were washed with 1X PBS (pH 7.4) before adding 50 uL of enzyme-free cell dissociation buffer (Thermo) to three wells and incubated for 5 min at room temperature. The cells were collected and spun down at 800xg for 8 min. Buffer was removed and 20 uL of RIPA Lysis and Extraction buffer (Thermo) supplemented with SIGMAFAST™ Protease Inhibitor Cocktail tablets (Sigma-Aldrich) were added to each well and incubated for 5 min on ice. The cell lysate was collected and centrifuged for 15 min at 14,000xg at 4°C. The supernatant was collected and combined with 4X SDS-PAGE Sample Loading Buffer and boiled at 95°C for 5 min. 30 uL of each transfection sample (along with non-transfected negative control) was run through a 4%–20% gradient Tris-Glycine gel (BioRad) and transferred to a PVDF membrane using the mixed molecular weight protocol of the Trans-Blot Turbo System (BioRad). The membrane was blocked with 5% milk in TBS-T (20 mM Tris-HCl pH 8.0, 150 mM NaCl, 0.05% Tween-20) at room temperature and with agitation. After 1 h, the HA polyclonal antibody (1:5000 dilution, Proteintech) was added and incubated overnight at 4°C with agitation. The membrane was washed three times with TBS-T and an IRDye 680 RD Goat anti-Rabbit IgG secondary antibody (1:50,000 dilution, LI-COR) was added and incubated for 1h at room temperature. The membrane was washed three times with TBS-T after which a LI-COR processor was used to develop the western blot. Membranes in figures were cropped to show area of interest.

### Biolayer interferometry assays

The BLI assays used to observe binding kinetics between EPCR and LayV G along with ephrinB2 and NiV G were performed on an Octet Red 96e (Sartorius). Octet AHC2 anti-human Fc biosensors (Sartorius) were prehydrated for 10 minutes in 10x Kinetics Buffer (Sartorius) which was also used as the running buffer of the entire experiment. Human-Fc tagged EPCR (ACROBiosystems, PRR-H5254) or ephrinB2-Fc were then loaded onto the biosensors to a 1 nm shift, baselined then dipped into a 1:3 serial dilution starting at 10 nM down to 1.37 pM of LayV G ectodomain tetramer or NiV G ectodomain tetramer for their respective ligand with the final well with only containing 10x Kinetics Buffer. Using Sartorius’ Octet Analysis 11.1 software the data was baseline subtracted and fitted using a 1:1 model before being replotted using Graphpad Prism 11.1. Each assay was run at 30 °C and 1000 RPM. We note that the tetrameric nature of LayV G and NiV G lead to marked binding avidity increasing the apparent affinities.

### Cryo-electron microscopy sample preparation, data collection, processing and model building

LayV G/EPCR complex was prepared by mixing a 2:1 molar ratio of the EPCR with LayV G ectodomain (oP4H-LayV G_SM_) before incubation for one hour on ice. Prior to freezing, a detergent mixture was added to 3 mg/ml LayV G/ EPCR complexes to make 0.0048% FOM, 0.06% Bril-35, 0.05% CHAPSO, 0.25% Amphipol 8-35 final concentration. For Cryo-EM grid preparation, 3 µL of the 3 or 1mg/ml of LayV G/EPCR complex with or without detergent samples were loaded onto freshly glow discharged R 2/2 UltrAuFoil (Quantifoil, Q130800) grids prior to plunge freezing using a Vitrobot Mark IV (ThermoFisher Scientific) with a blot force of −2 and 2 second blot time at 100% humidity and 4°C or 22°C temperature. Data were acquired using a TFS Glacios transmission electron microscope operated at 200 kV and equipped with a Gatan K3 Summit direct electron detector. Automated data collection was carried out using Leginon(Suloway et al. 2005)at a nominal magnification of 45,000x, a pixel size of 0.89 Å, a dose rate of 7.5 counts/pixel/second with each movie acquired in counting mode fractionated in 100 frames of 50 ms. A total 10,241 movies were collected with a defocus range between −0.2 and −3 μm.

Movie frame alignment, estimation of the microscope contrast-transfer function (CTF) parameters, initial particle picking, and extraction were carried out using Warp(v1.1.0)(Tegunov and Cramer 2019). To select well-defined particle images, reference-free 2D classification followed by Ab initio 3D reconstruction and heterogeneous 3D refinement were performed in CryoSPARC (v5.0.6)(Punjani et al. 2017). 3D refinement was carried out using non-uniform refinement (NUR) with per-particle defocus refinement with C4 symmetry in CryoSPARC (v5.0.6). To minimize the potential loss of rare views of particles, the second rounds of particle picking were carried out using Template based particle picking and Topaz with training using Warp-picked particle data set belonging to the selected best 2D classes or 3D refinement model from Warp data set followed by particle extraction, 2D classifications, heterogeneous 3D refinement and NUR with per-particle defocus refinement with C4 symmetry in CryoSPARC (v5.0.6). After removal of duplicated particles from each data sets, the combined particle stacks from the Warp, Topaz and Template based picking runs were processed together in the particle extraction with recentering, junk particles removed around the gold/carbon surface by micrograph junk detention and 3D non-uniform refinement with per-particle defocus refinement. Next, bayesian polishing was performed in Relion (v5.0)(Zivanov et al. 2018) using unbinned particle stacks, with data export assisted by pyem (v0.5) (https://github.com/asarnow/pyem), followed by 30 beam tilt group clusterings using K-means. During bayesian polishing in Relion, particles were re-extracted with a box size of 448 pixels and a pixel size of 1.0 Å. To select well-defined polished particles, reference-free 2D classification and five round heterogeneous 3D refinements were performed in CryoSPARC (v5.0.6). The final 3D refinement was carried out using non-uniform refinement with per-particle defocus refinement, global CTF refinement and C4 symmetry in CryoSPARC (v5.0.6), yielding a map with a global resolution of 2.9 Å. To further improve the Cryo-EM maps, local refinement with a global mask at the center of Head tetramer or a soft local mask comprising LayV G Head/EPCR domains was conducted using CryoSPARC (v5.0.6), resulting in a 2.92 Å and 2.91 Å resolution reconstruction, respectively. Final reconstruction statistics are summarized in Table1. Reported resolutions are based on the 0.143 gold-standard Fourier shell correlation (FSC) criterion and Fourier shell correlation curves were corrected for the effects of soft masking by high-resolution noise substitution (Rosenthal and Henderson 2003; Chen et al. 2013). Local resolution estimation, filtering, and sharpening were carried out using CryoSPARC (v5.0.6).

For model building, we started from the previously determined LayV G (PDB 8TVI) and EPCR (PDB 7OKS) structures which we docked into density. Manual rebuilding and real-space refinement with unsharpened and sharpened maps were carried out in UCSF ChimeraX (v1.12)(Pettersen et al. 2004)and Coot (v0.9.8.95EL)(Emsley et al. 2010). The final LayV G/EPCR complex model was refined using Phenix (v2.2.1)(Liebschner et al. 2019) and validated using MolProbity (Chen et al. 2010).

---

**Figure S1.**
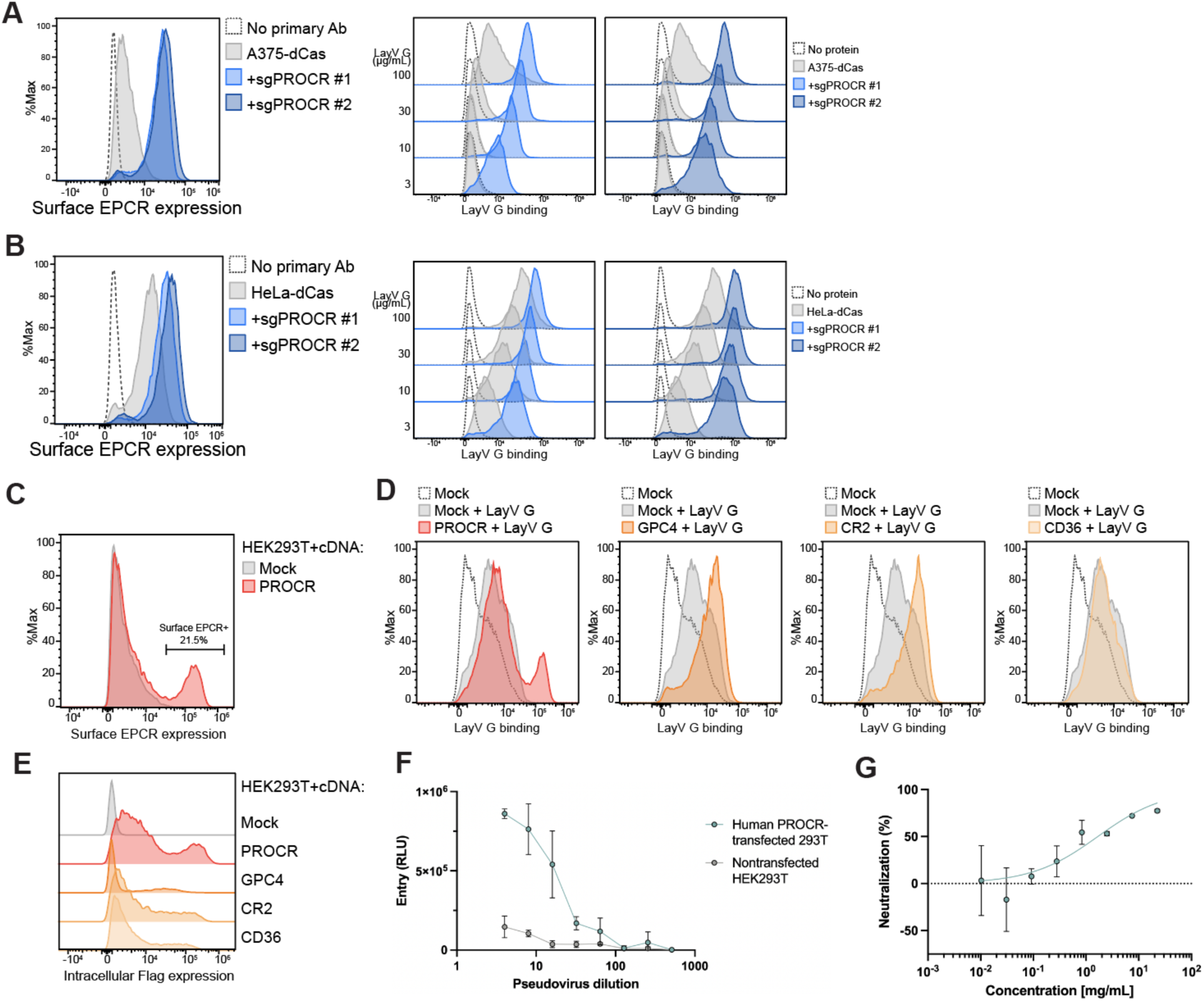
Validation of the role of EPCR in Langya pseudovirus entry into cells. **A-B.** Flow cytometry analysis of binding of the LayV G ectodomain tetramer to A375 (A) or Hela (B) cells transduced with activating sgRNAs targeting *PROCR* to induce EPCR expression. (Left) Quantification of cell surface EPCR expression using a rat monoclonal anti-human EPCR antibody (clone RCR-401). (Right) Titration of LayV G binding at various concentrations. **C.** Surface expression of EPCR in HEK293T cells transfected with PROCR cDNA or empty vector (mock).**D-E.** Flow cytometry analysis of LayV G ectodomain tetramer binding to HEK293T cells ectopically expressing the four top hits from the screen, PROCR, GPC4, CR2, and CD36 (D), and intracellular staining of the C-terminal Flag epitope tag to confirm expression of each construct (E). **F.** Entry of various dilutions of non-replicating, lentiviral particles pseudotyped with the LayV F/G glycoproteins in HEK293T cells transiently transfected or not with human EPCR. Each data point represents the average of three technical replicates. One biological replicate is shown, with error bars showing standard deviation. **G.** Soluble EPCR-mediated neutralization of lentiviral particles pseudotyped with the LayV F and G glycoproteins. Each data point represents the average of two technical replicates. The second biological replicate is shown (first one shown in Figure 1F), with error bars showing standard deviation.

**Figure S2.**
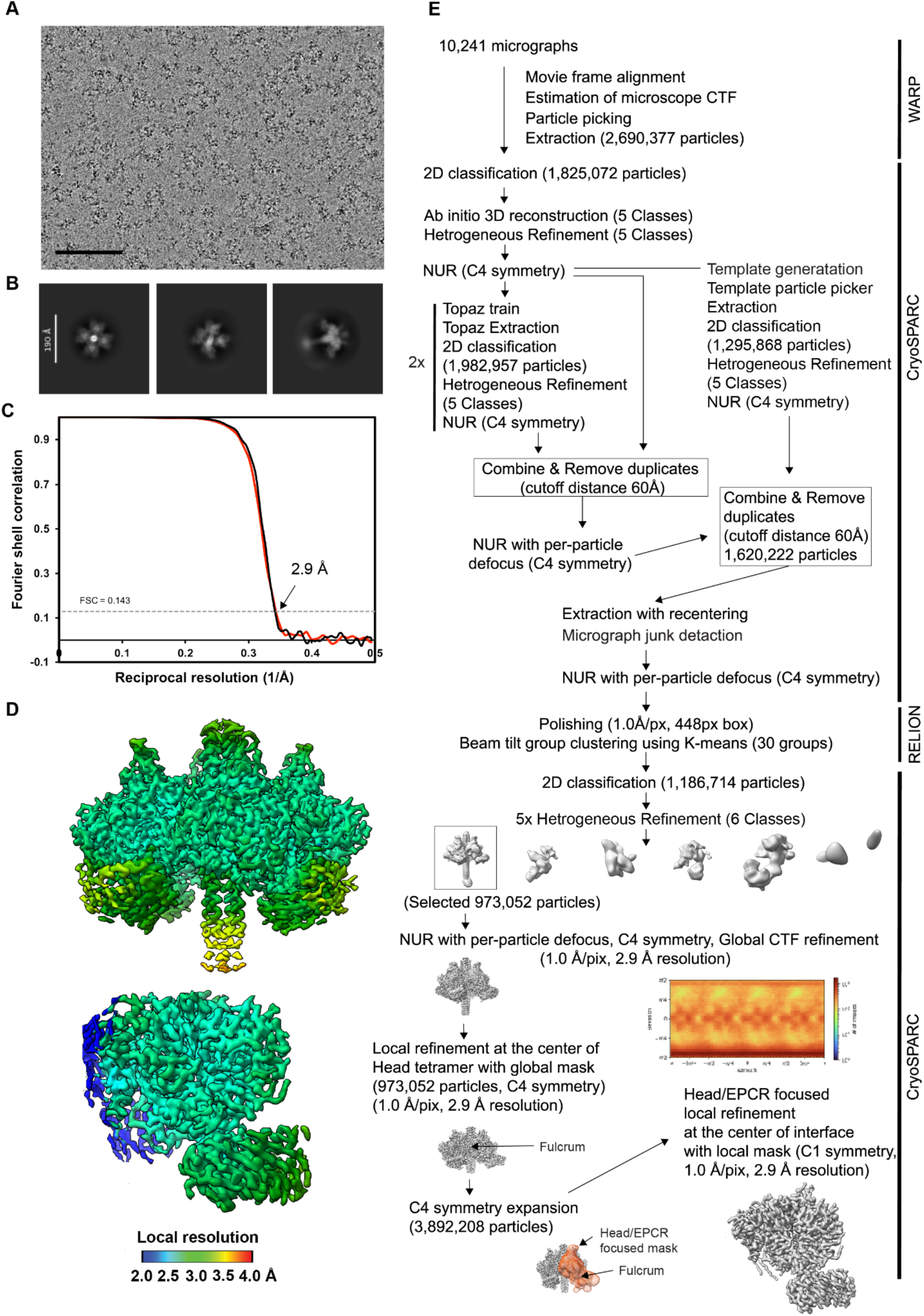
Cryo-EM data processing of the human EPCR-bound LayV G dataset. **A-B,** Representative electron micrograph and 2D class averages of the human EPCR ectodomain bound to the LayV G ectodomain (oP4H-LayV G_SM_) tetramer complex embedded in vitreous ice. Scale bars: 100 nm (A) and 190 Å (B). **C**, Gold-standard Fourier shell correlation curves for the global reconstruction (black) and for the reconstruction obtained by local refinement of the G head domain and EPCR (red). The 0.143 cutoff is indicated by a horizontal dashed line. **D**, Local resolution estimation calculated using cryoSPARC and plotted on the sharpened global (top) and local (bottom) refinement maps. **E**, Data processing flowchart. CTF: contrast transfer function; NUR: non-uniform refinement. The angular distribution of particle images calculated using cryoSPARC is shown as a heat map.

**Fig S3.**
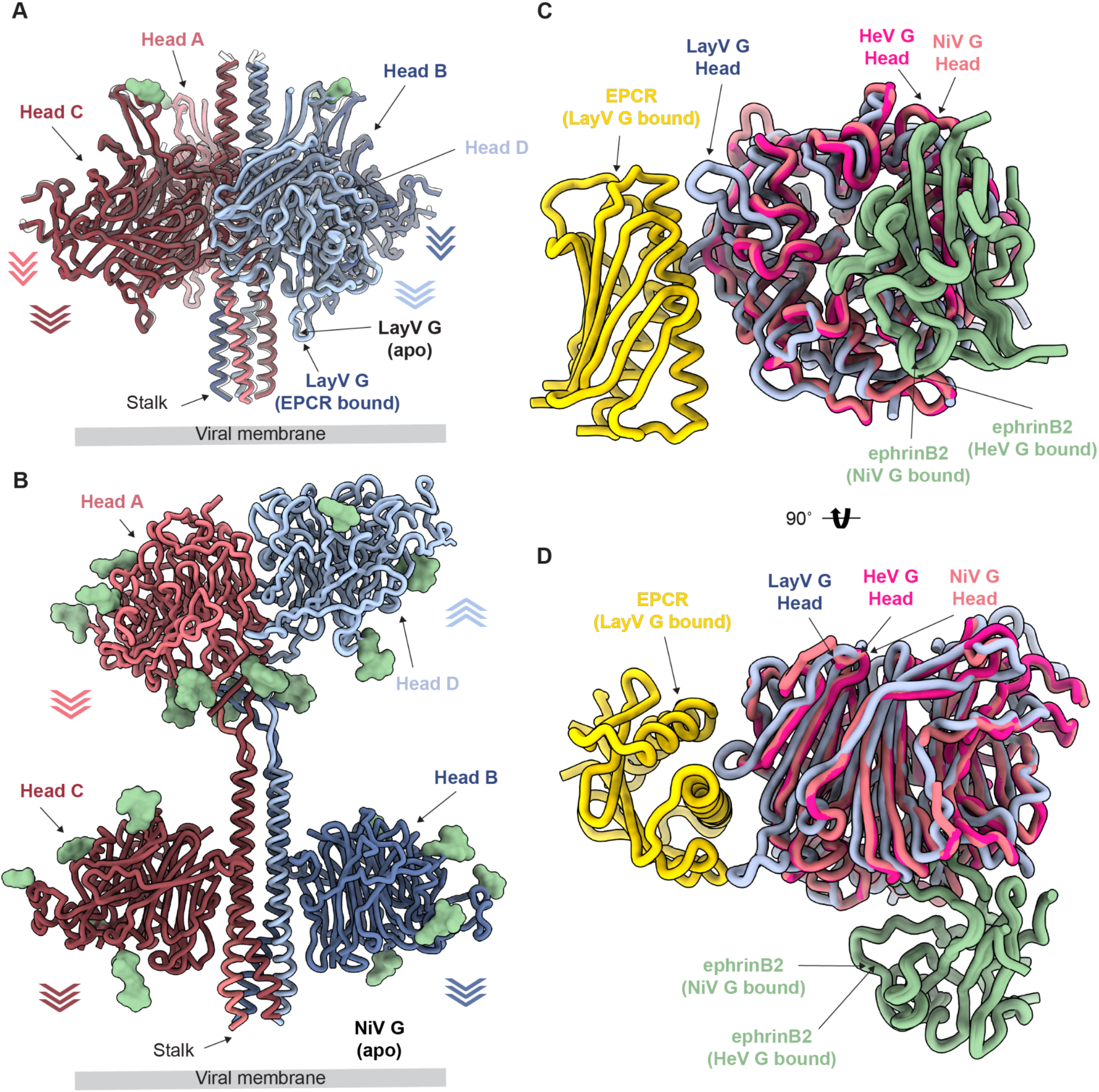
Architecture of the LayV G ectodomain tetramer and comparison of the receptor binding sites for LayV G and NiV G. **A,** Superimposition of the global cryoEM structure of the EPCR-bound LayV G tetramer and of the apo LayV G tetramer (PDB 8TVI, grey). EPCR is omitted for clarity. **B,** CryoEM structure of the NiV G tetramer (PDBs 7TXZ and 7TY0) with the bound nAH1.3 Fabs omitted for clarity. In panels A-B, N-linked glycosylation sites are rendered as green surfaces and arrows indicate the orientation of each head domain. **C-D**, Superimposition of the local cryoEM structure of the EPCR-bound LayV G head domain (yellow and blue, respectively) with the crystal structure of the ephrinB2-bound NiV G head domain (green and light pink, respectively, PDB 2VSM) and the crystal structure of the ephrinB2-bound HeV G head domain (green and light red, respectively, PDB 2VSK) shown in two orthogonal orientations.

**Figure S4.**
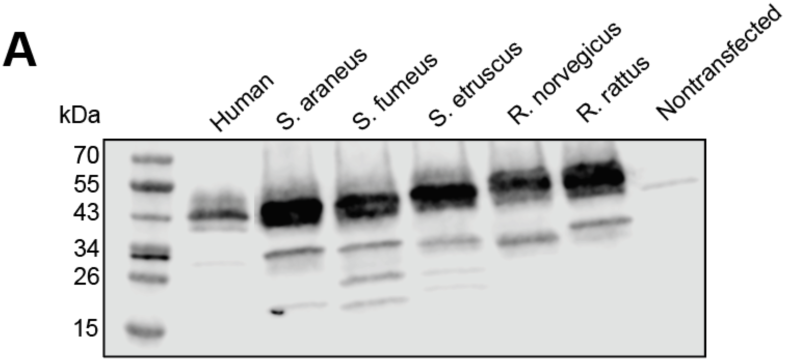
Langya and Mojiang pseudovirus entry and western blot of EPCR orthologs. **A-B.** Entry of various dilutions of non-replicating, lentiviral particles pseudotyped with the LayV or MojV F/G glycoproteins in HEK293T cells transiently transfected or not with the indicated EPCR orthologs. Each data point represents the average of two technical replicates. One biological replicate is shown per panel (three in total), and error bars show the standard deviation. **C.** Western blot of EPCR orthologs transiently transfected into HEK293T cells using anti-HA tag as primary antibody.

**Table S1.** *Surfaceome* CRISPRa screening results.

**Table S2.** Cryo-EM data collection and refinement statistics.

|  | **LayV G - human EPCR**<br>**(global refinement)**<br>PDB ###<br>EMD-### | **LayV G - human EPCR**<br>**(Head/EPCR focused local refinement)**<br>PDB ###<br>EMD-### |
|---|---|---|
| **Data collection and processing** |  |  |
| Magnification | 45,000 | 45,000 |
| Voltage (kV) | 200 | 200 |
| Electron exposure (e–/Å<sup>2</sup>) | 47 | 47 |
| Defocus range (µm) | -0.2 - -3.0 | -0.2 - -3.0 |
| Pixel size (Å) | 0.89 | 0.89 |
| Symmetry imposed | C4 | C1 |
| Initial particle images (no.) | 2,690,377 | 2,690,377 |
| Final particle images (no.) | 973,052 | 3,892,208 |
| Map resolution (Å) | 2.9 | 2.9 |
| FSC threshold | 0.143 | 0.143 |
| **Refinement** |  |  |
| Model resolution (Å) | 3.0 | 3.0 |
| FSC threshold | 0.5 | 0.5 |
| Map sharpening *B* factor (Å<sup>2</sup>) | -121 | -121 |
| Model composition |  |  |
| Non-hydrogen atoms | 2,1116 | 4,750 |
| Protein residues | 2,704 | 603 |
| Ligands | 32 | 8 |
| Water | 0 | 0 |
| *B* factors (Å<sup>2</sup>) |  |  |
| Protein | 33.3 | 33.8 |
| Ligand | 69.7 | 74.1 |
| Water |  |  |
| R.m.s. deviations |  |  |
| Bond lengths (Å) | 0.002 | 0.003 |
| Bond angles (°) | 0.710 | 0.584 |
| **Validation** |  |  |
| MolProbity score | 1.30 | 1.63 |
| Clashscore | 3.05 | 5.21 |
| Poor rotamers (%) | 0.62 | 1.49 |
| Ramachandran plot |  |  |
| Favored (%) | 96.70 | 94.62 |
| Allowed (%) | 3.15 | 5.21 |
| Disallowed (%) | 0.15 | 0.17 |

